# Na/K-ATPase Signaling in Adipocytes Promotes Atherosclerosis

**DOI:** 10.1101/2025.08.18.670994

**Authors:** Bruno S. Goncalves, Yaxin Wang, Sneha S. Pillai, Hari V. Lakhani, Aslam Chaudhry, Joseph I. Shapiro, Roy L. Silverstein, Komal Sodhi, Yiliang Chen

## Abstract

**BACKGROUND:** Adipocyte dysfunction is closely associated with oxidative stress and chronic inflammation, which contribute to systemic metabolic disturbances and atherosclerosis. We previously identified Na/K-ATPase (NKA) α1 as a signal transducer that activates Src family kinases and promotes oxidative stress and inflammation in various cell types, including adipocytes and macrophages. NaKtide, a peptide inhibitor of NKA signaling, has been shown to reduce systemic oxidative stress and inflammation *in vivo*. Here, we investigated the role of adipocyte-specific NKA signaling in atherosclerosis.

**METHODS:** Adipocyte-specific NaKtide (Adipo-NaKtide) was delivered to *Apoe*^-/-^ mice using a lentiviral vector under the adiponectin promoter. The mice were then fed a Western Diet (WD) for 12 weeks to induce atherosclerosis and then assessed for atherosclerotic plaque burden the aortic arch and at the level of the aortic sinus. Inflammatory and oxidative stress markers were analyzed in adipose tissue and plasma.

**RESULTS:** Adipo-NaKtide reduced atherosclerotic plaque area by 67% in the aortic arch and 47% in the aortic sinus. CD68^+^ macrophage and α-SMA^+^ smooth muscle cell content in the aortic sinus were decreased by 45% and 53%, respectively. These vascular improvements were accompanied by dampened adipose tissue inflammation and oxidative stress, improved glucose tolerance, and lowered systemic inflammation.

**CONCLUSIONS:** These findings highlight a critical contributing role for adipocyte NKA signaling in atherosclerosis, suggesting an important endocrine and/or paracrine influence of adipose tissue on large artery atherogenesis, and supporting the therapeutic potential of targeting NKA in cardiometabolic disease.

## INTRODUCTION

Atherosclerosis is a chronic inflammatory condition, driven by lipid accumulation in the arterial wall, systemic metabolic dysfunction, and oxidative stress^1–3^. Among the metabolic organs contributing to this pathological state, adipose tissue plays a pivotal role^4,5^. Under conditions such as obesity or high-fat feeding, adipocytes undergo functional changes that result in increased secretion of pro-inflammatory cytokines and reactive oxygen species (ROS), which in turn exacerbate systemic inflammation and promote vascular injury^6,7^.

The Na/K-ATPase (NKA) α1 subunit, beyond its canonical cation transporter function, was identified nearly two decades ago as a signal transducer through its ability to interact directly with Src family kinases (SFK)^8–11^. The cytoplasmic domain of NKA α1 sequesters SFKs at the inner leaflet of the plasma membrane, holding them in an inactive, inaccessible state. In response to extracellular stimuli, conformational changes in NKA release activated SFKs triggering downstream signaling cascades that promote oxidative stress and inflammation in multiple cell types, including macrophages and adipocytes^9,11^. A 20-amino acid mimetic peptide based on the sequence of the SFK-binding domain of the NKA α1 subunit was developed^12^ to inhibit the NKA-SFK signaling axis, and termed NaKtide. NaKtide binds to the conserved SFK kinase domain, effectively blocking its activation and downstream signaling. Our previous studies showed that systemic administration of NaKtide reduced oxidative stress, improved metabolic parameters, and attenuated inflammation in animal models of chronic diseases^13–15^. However, the specific role of adipocyte NKA signaling in the progression of atherosclerosis has not yet been defined.

We previously demonstrated that NKA α1 forms a functional complex with CD36^16^ in adipocytes. CD36 is a fatty acid transporter and scavenger receptor known to play an important role in macrophage uptake of oxidized low-density lipoproteins (oxLDL), foam cell formation, and atherosclerosis^17^. The NKA-CD36 signaling axis promotes lipid accumulation, ROS generation, pro-inflammatory cytokine expression, and metabolic reprogramming under atherogenic conditions in various cell types, including adipocytes^16,18,19^. These proatherogenic effects were abolished by pNaKtide, a cell-permeable version of NaKtide *in vitro*^16^, suggesting that NKA signaling may contribute to vascular inflammation and atherogenesis. Despite these findings, the *in vivo* significance of adipocyte-specific NKA signaling in atherosclerosis remains unknown. In this study, we addressed this gap by expressing NaKtide selectively in adipocytes using a lentiviral vector under the control of the adiponectin promoter (Adipo-NaKtide) in high fat diet-fed *Apoe*^-/-^ mice and found that it reduced atherosclerotic burden in the aortic arch and sinus, dampened adipose tissue inflammation and oxidative stress, improved glucose tolerance, and lowered circulating levels of systemic inflammatory markers. These findings reveal a previously unrecognized role of adipocyte NKA signaling in the pathogenesis of atherosclerosis and suggest that targeting adipocyte NKA could be an effective therapeutic approach to limit metabolic inflammation in cardiovascular disease.

## MATERIALS AND METHODS

### Mouse Models and *In Vivo* Studies

All animal studies were approved by the Marshall University and Medical College of Wisconsin Animal Care and Use Committees in accordance with the National Institutes of Health (NIH) Guide for the Care and Use of Laboratory Animals. ApoE knockout mice (*Apoe*^-/-^) (6-8 weeks old, male) were purchased from Jackson Laboratory. Upon arrival at the Robert C. Byrd Biotechnology Science Center Animal Resource Facility (ARF), the mice were housed in designated rooms equipped with cages that supplied purified air under a 12-hour light/dark cycle. Mice were fed a normal chow diet or a high-fat Western Diet (WD) for 3 months. WD is a well-known model of diet-induced oxidative stress and metabolic syndrome, and it contributes to chronic metabolic imbalance and the development of atherosclerosis^19,20^. The WD was purchased from Envigo (Indianapolis, IN) and contained 42% fat, 42.7% carbohydrate, and 15.2% protein yielding 4.5 KJ/g.

Lentiviral vectors expressing either green fluorescent protein (GFP) tagged NaKtide or GFP alone under the control of an adiponectin promoter were constructed by VectorBuilder Inc. to yield adipocyte-specific expression as previously reported ^21^. Lentivirus constructs (100 µl, 2×10^9^ TU/ml) in saline were intraperitoneally (IP) injected into *Apoe*^-/-^ mice as described previously^21^. Mice were euthanized 12 weeks after the dietary interventions and at the time of termination, blood, visceral fat (VFAT), heart, and aorta were collected. Plasma was isolated, flash-frozen in liquid nitrogen, and maintained at -80 °C until measurement of plasma biomarkers. Mice were euthanized by ketamine overdose (100 mg/Kg body weight) and perfused via the left ventricle with 10 ml PBS. After perfusion, the aorta was dissected, cleaned, and fixed in 4% paraformaldehyde.

Heart tissues containing the aortic roots and adipose tissues were carefully dissected and embedded in optimal cutting temperature (OCT) solution for cryosection. Some heart, aorta and adipose tissues were flash-frozen in liquid nitrogen and maintained at -80 °C for Western blots and quantitative real-time PCR.

### Measurement of Protein Carbonylation

Protein carbonylation was measured in VFAT, using a Protein Carbonyl ELISA Assay Kit (BioCell Corporation, Auckland, New Zealand), according to the manufacturer’s protocol.

### Estimation of SFK active site phosphorylation [pY419] by Western Blot analysis

VFAT was homogenized with RIPA buffer, and pSFK was detected with a polyclonal anti-SFK [pY419] phospho-specific antibody (Invitrogen). The same membranes were then stripped and blotted with a monoclonal antibody against total SFK (Santa Cruz). Expression of pSFK was expressed as the ratio of pSFK/tSFK, with measurements normalized to control samples.

### RNA extraction and real-time PCR

Total RNA was extracted from snap-frozen mouse VFAT and aorta tissue using RNeasy Protect Mini Kit (QIAGEN), according to the manufacturer’s instructions. Glyceraldehyde- 3-phosphate dehydrogenase (GAPDH) was used as an endogenous control. Specific predesigned mouse-specific primers (IDT DNA Technologies) were used for *Cd36*, interleukin-6 (*Il6*), tumor necrosis factor-alpha (*Tnf*), peroxisome proliferator-activated receptor gamma coactivator 1-alpha (*Ppargc1a*), monocyte chemoattractant protein-1 (*Ccl2*), triggering receptor expressed on myeloid cells 2 (*Trem2*), pyruvate kinase muscle (*Pkm*) and F4/80 (*Adgre1*). For microRNA (miR) extraction from plasma samples, miRNeasy Serum/Plasma Kits (Qiagen) were used according to the manufacturer’s protocol. miRCURY LNA RT Kits (Qiagen) were used to synthesize cDNA. Mouse-specific miR primers were used for miR-27b-3p, and miR-34a-3p (Qiagen). The respective primers were combined with miRCURY LNA SYBR® Green PCR Master Mix (Qiagen) at a final volume of 10 μl to perform RT-PCR reactions in a 96-well plate. miR expressions of all experimental samples were normalized to a mouse U6 amplicon. For both mRNA and miR, two technical replicates were used per sample to ensure the accuracy of the RT-PCR amplification data, which was run on a 7500 Fast Real-Time PCR System (Applied Biosystems). The comparative threshold cycle (C_t_) method was used to calculate the fold amplification, as specified by the manufacturer.

### Measurement of plasma cytokines and glucose profile

Plasma IL-6 and TNF-α were measured using Enzyme-Linked Immunosorbent Assay (ELISA) kits, according to the manufacturer’s protocol (Abcam, Cambridge, MA). We also performed the glucose tolerance test (GTT) using a glucometer. After the fasting period, a 10% glucose solution (2 g/kg body weight) was IP injected. Samples were taken from the tail vein at 0, 30, 60, 90, and 120 minutes after glucose injection.

### Oil Red O staining of the aortic arch and aortic sinus

To quantify the atherosclerotic plaque area, aortic arches were isolated and stained with Oil Red O (ORO). Briefly, aortas were opened longitudinally and were rinsed with PBS, followed by staining with 2.4% ORO solution for 15 minutes. Then, tissues were destained in 60% isopropanol until the background was clear. The lesion area was quantified as a percentage of the ORO–stained area in the total aorta arches area using ImageJ software.

For aortic sinus staining, 6μm serial cross-sections of aortic root tissue were prepared and stained with 2.4% ORO solution for 30 minutes, followed by washing in PBS three times. The cross-sectional lesion area for each mouse was calculated using the mean value of 3-4 series of sections. The lesion area was quantified as a percentage of the ORO–stained area in the total aorta sinus area using ImageJ software.

### Immunofluorescence assay

Immunofluorescence studies were performed on OCT-embedded cryosections (6μm) of adipose and heart tissue. The cryosections were fixed with 4% paraformaldehyde at room temperature (RT) for 15 minutes, washed, and blocked with PBS containing 5% bovine serum albumin at RT for 2 hours. The sections were subsequently incubated overnight with primary NaKtide antibody (dilution of 1:30) at 4°C, followed by washing and probing with secondary antibody (1:1000, Alexa Fluor 647 Red), and the slides were then mounted with DAPI. Expression of GFP and NaKtide was determined using GFP and RFP filters, respectively, on a Nikon Eclipse 80i microscope equipped with a Nikon camera head DS-Fi1 (Nikon).

For immunofluorescence studies on the aortic sinus, after the blocking step, tissue sections were incubated overnight with either primary CD68 antibody (Bio-Rad, MCA1957GA, dilution of 1:100) or primary α-smooth muscle actin (α-SMA) antibody (Invitrogen, 14-9760-82, dilution of 1:100) at 4°C overnight. The following day, the slides were washed three times with PBS and stained with secondary antibody Alexa Fluor 488 (Invitrogen, dilution of 1:200) for 1 hour at RT. After PBS washing three times, the slides were visualized using an EVOS 7000 microscope (Thermo Scientific). Results were expressed as a percentage of cell marker-positive areas among the total aortic sinus area.

### Statistical analysis

Results are presented as means ± standard error (SEM). Comparisons between the groups were performed using a one-way and two-way analysis of variance (ANOVA), followed by Tukey’s test comparison for post-hoc analysis. Statistical significance was assigned at p<0.05. Data analyses were performed using GraphPad Prism 10.3 (GraphPad Software, San Diego, CA).

## RESULTS

### Adipocyte-specific NaKtide expression attenuated atherosclerotic plaque formation in *Apoe*^-/-^ mice fed WD

Tissue specificity of the lentiviral construct expressing GFP-NaKtide under the control of the adiponectin promoter (Adipo-NaKtide) was assessed by immunofluorescence microscopy on visceral fat (VFAT) from *Apoe*^-/-^ mice using an anti-NaKtide antibody. In mice injected with the Adipo-NaKtide construct, there was robust co-localization of GFP (green) and NaKtide (red) signals in VFAT (Figure 1A. In contrast, no NaKtide signal was detected in VFAT from mice injected with the control lentiviral vector (Adipo-GFP) or in heart tissues from either vector treatment group (Figure 1A and Suppl. Figure 1A).

**Figure 1.**
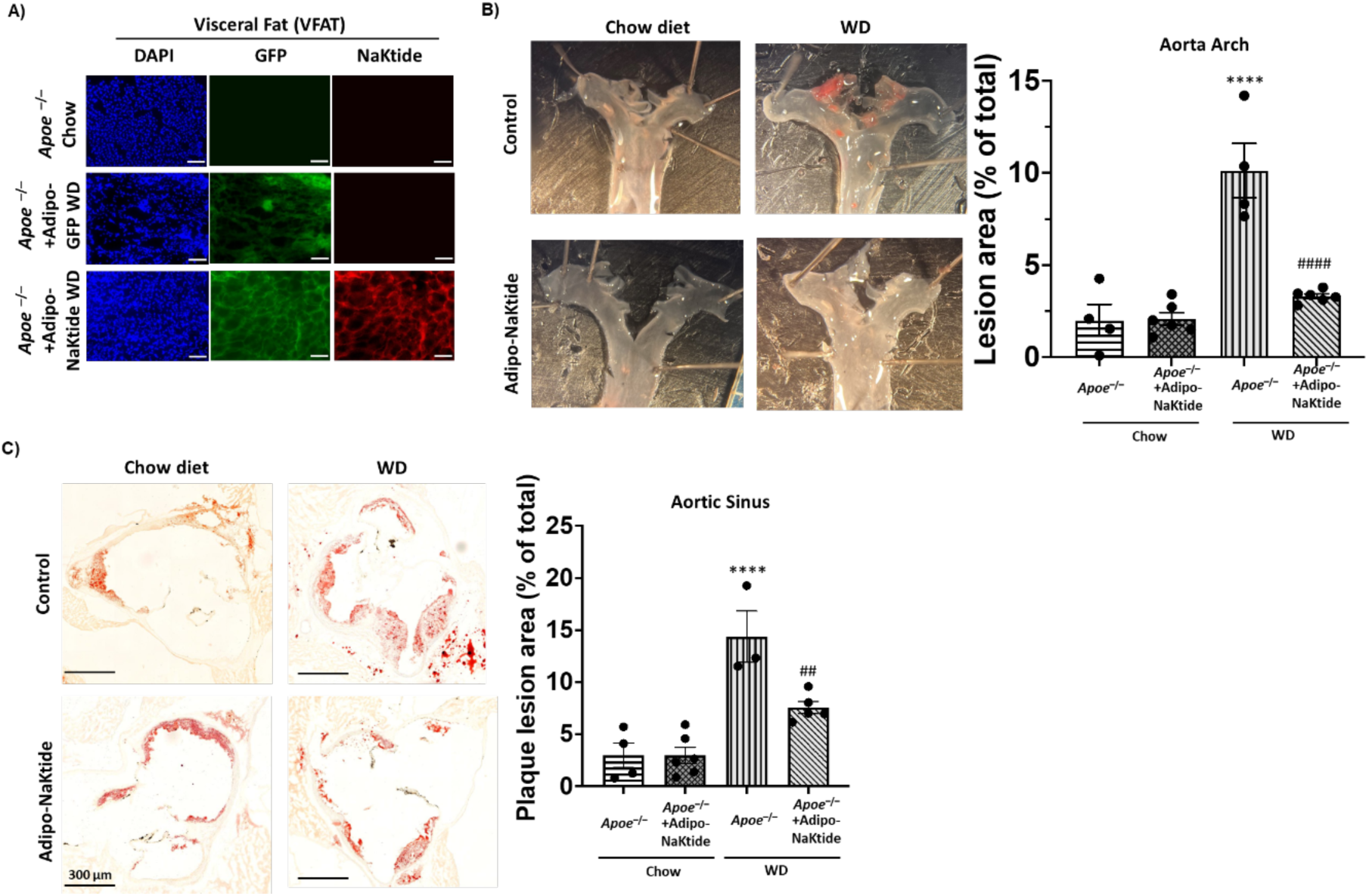
Adipocyte-specific expression of NaKtide attenuates atherosclerotic plaque formation in *Apoe*^-/-^ mice fed a WD. (**A**) Immunofluorescence staining demonstrating expression of GFP (green) and NaKtide (red) in visceral adipose tissue of mice transfected with Adipo-NaKtide (N=6/group). (**B**) Left panel: representative ORO- stained images of aortic arch from *Apoe*^-/-^ mice fed with chow diet or WD for 12 weeks in the absence or presence of Adipo-NaKtide expression. Right panel: ORO-positive areas were quantified and shown in the bar graph (N=4-6/group). (**C**) Left panel: representative ORO-stained images of aortic sinus from *Apoe*^-/-^ mice fed with chow diet or WD for 12 weeks in the absence or presence of Adipo-NaKtide expression. Right panel: ORO-positive areas were quantified and shown in the bar graph (N=3-6/group). Scale bar: 300 μm. Values represent means ± SEM. ****p<0.0001 vs. *Apoe*^-/-^ chow; ^##^p<0.01 vs. *Apoe*^-/-^ WD; ^####^p<0.0001 vs. *Apoe*^-/-^ WD.

After 12 weeks of WD, atherosclerotic plaque formation in the aortic arch of *Apoe*^-/-^ mice with or without Adipo-NaKtide was evaluated by *en face* ORO staining. WD-fed mice had lipid-rich atherosclerotic plaques covering 10.12±1.70% of total arch area, compared to 1.99±1.00% in the chow-fed mice. Adipo-NaKtide treatment reduced plaque area in the aortic arch by 67% in the WD-fed mice (p<0.0001) (Figure 1B). Similarly, cross sectional plaque area at the level of the aortic sinus was reduced by 48% (p<0.01) in WD-fed mice treated with Adipo-NaKtide lentivirus compared to the chow diet control (14.37±3.01% vs 7.55±0.66% plaque area) (Figure 1C). Since expression of the control lentiviral vector (Adipo-GFP) did not affect atherosclerotic plaque development in *Apoe*^-/-^ mice (Supplementary Figure 1B), all subsequent experiments were conducted using the same four groups shown in Figure 1B.

### Adipocyte-specific NaKtide expression reduced macrophage infiltration and smooth muscle cell content in aortic sinus lesions of *Apoe*^-/-^ mice fed WD

Macrophages play a central role in the development and progression of atherosclerosis^22^. Immunostaining for the macrophage marker CD68 revealed that Adipo-NaKtide expression reduced CD68-positive areas in the aortic sinus of WD-fed mice by 45% from 7.88±0.39% to 4.37±0.86% (Figure 2A). In addition to macrophage accumulation, smooth muscle cell (SMC) proliferation contributes to plaque growth and stability^23,24^. Immunostaining for α-smooth muscle actin (α-SMA) showed a 53% reduction in α-SMA-positive areas in the aortic sinus (p<0.001) from 6.59±0.69% to 3.11±0.58% in the Adipo-NaKtide treated WD-fed mice (Figure 2B).

**Figure 2.**
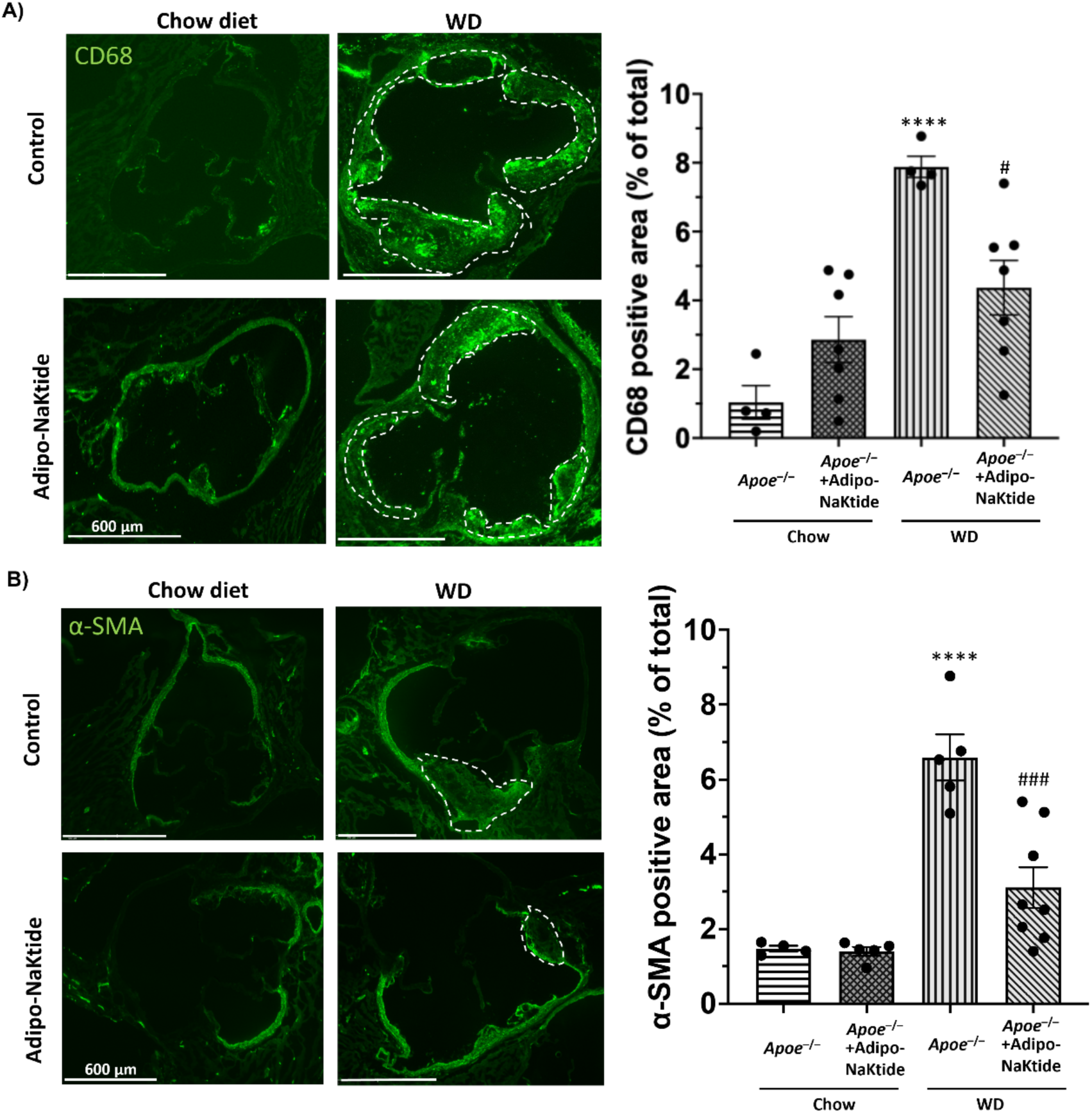
Adipocyte-specific expression of NaKtide reduces the CD68 and α-SMA-positive area in *Apoe*^-/-^ mice fed a WD. (**A**) Left panel: representative fluorescence images of aortic sinus immunostained with anti-CD68 (green) from *Apoe*^-/-^ mice fed with chow diet or WD for 12 weeks in the absence or presence of Adipo-NaKtide expression. Right panel: CD68-positive areas were quantified and shown in the bar graph. Scale bar: 600 μm (N=4-7/group). (**B**) Left panel: representative fluorescence images of aortic sinus immunostained with anti-α-SMA (green) from *Apoe*^-/-^ mice fed with chow diet or WD for 12 weeks in the absence or presence of Adipo-NaKtide expression. Right panel: α-SMA-positive areas were quantified and shown in the bar graph. Scale bar: 600 μm (N=4-8/group). Values represent means ± SEM. ****p<0.0001 vs. *Apoe*^-/-^ chow; ^#^p<0.05 vs. *Apoe*^-/-^ WD, ^###^p<0.001 vs. *Apoe*^-/-^ WD.

### Adipocyte-specific NaKtide expression suppressed WD-induced inflammation in aorta of *Apoe*^-/-^ mice fed WD

mRNA levels of pro-inflammatory proteins *Tnf*, *Il6*, and *Ccl2* in aorta were increased by 3-5 fold in WD-fed *Apoe*^-/-^ mice compared to those fed chow diet (Figure 3A-C; p<0.01). Treatment with Adipo-NaKtide lentivirus downregulated expression of these genes in WD-fed *Apoe*^-/-^ mice by 39-44% compared to control animals (p<0.05). *Trem2* and *Adgre1* are markers of aortic lipid-laden foamy macrophages^25^ and *Pkm* encodes a glycolytic enzyme that is associated with abnormal glucose metabolism and chronic inflammation in macrophages from human patients with coronary artery disease^26^. Expression of these genes was upregulated by WD (between 153 and 320% increase, p<0.05, 0.001, and 0.01, respectively), and this was suppressed by treatment with Adipo-NaKtide lentivirus (Figure 3D-F). These data together show that adipose expression of NaKtide protected *Apoe*^-/-^ mice from diet-induced arterial inflammation and atherosclerosis.

**Figure 3.**
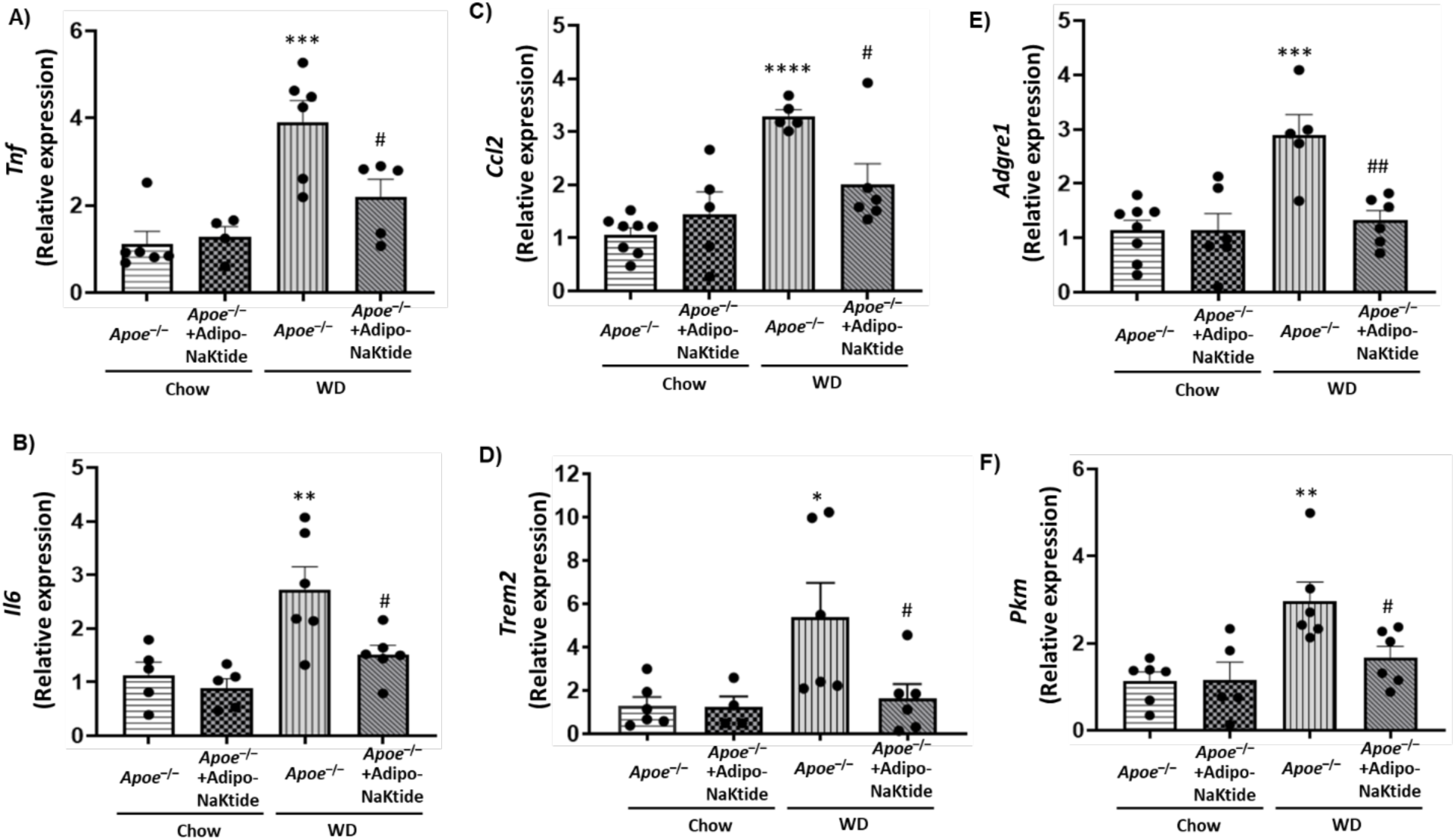
Adipocyte-specific expression of NaKtide improves inflammation and macrophage activation in aorta tissue in *Apoe*^-/-^ mice fed a WD. Quantitative real-time polymerase chain reaction (qRT-PCR) analysis for (**A**) *Tnf* (N=4-6/group); (**B**) *Il6* (N=5-6/group); (**C**) *Ccl2* (N= 5-8/group); (**D**) *Trem2* (N=4-6/group); (**E**) *Adgre1* (N=5-8/group); (**F**) *Pkm* (N=5-6/group). For all qRT-PCR GAPDH was used as a housekeeping gene. Values represent means ± SEM. *p<0.05 vs. *Apoe*^-/-^ chow; **p<0.01 vs. *Apoe*^-/-^ chow; ***p<0.001 vs. *Apoe*^-/-^ chow; ****p<0.0001 vs. *Apoe*^-/-^ chow; ^#^p<0.05 vs. *Apoe*^-/-^ WD; ^##^p<0.01 vs. *Apoe*^-/-^ WD.

### Adipocyte-specific NaKtide expression inhibited NKA signaling and reduced oxidative stress and inflammation in adipose tissue of *Apoe*^-/-^ mice fed WD

Adipose tissue levels of protein carbonylation, an indicator of oxidative stress, were increased 3.3-fold in WD-fed *Apoe^-/-^* mice compared to chow-fed animals (0.85±0.07 nmol/mg vs 0.26±0.02 nmol/mg) (Figure 4A). Adipo-NaKtide expression attenuated carbonylation levels to 0.45±0.03 nmol/mg (p<0.0001). Similarly, SFK phosphorylation was increased by 70±22% by WD, and this increase was abolished by Adipo-NaKtide expression (Figure 4B). Furthermore, Adipo-NaKtide expression also significantly attenuated WD-induced increased expression of genes that mediate adipocyte inflammation, including *Cd36*, *Il6*, and *Tnf (*Figure 4C, D and E). Adipo-NaKtide treatment also increased expression of *Ppargc1a*, a key regulator of mitochondrial biogenesis and oxidative stress^27^ (Figure 4F).

**Figure 4.**
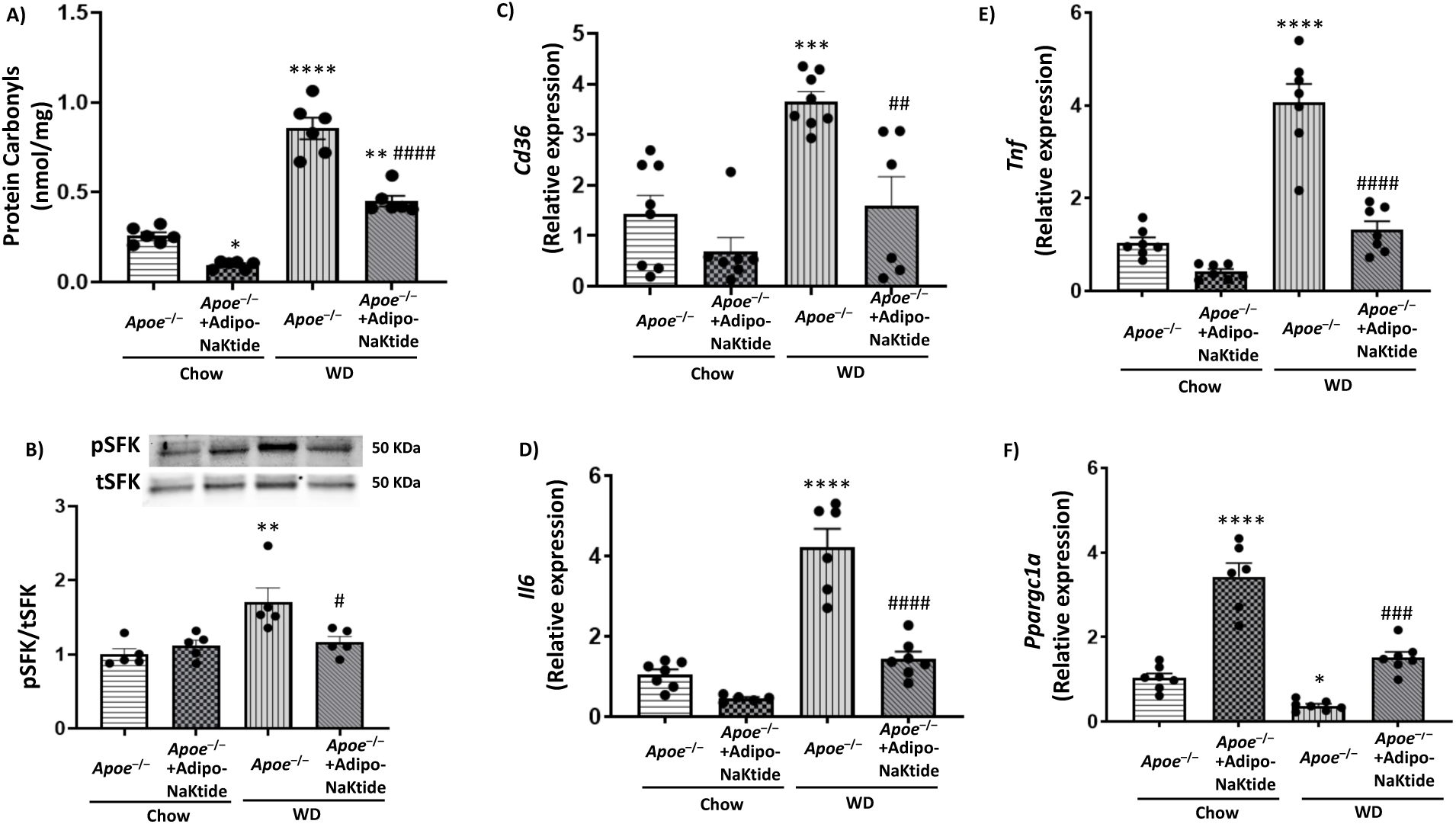
Adipocyte-specific expression of NaKtide improves oxidative stress, Na/K-ATPase signaling, and inflammation in VFAT in *Apoe*^-/-^ mice fed a WD. (**A**) Protein carbonylation (N=6/group). (**B**) pSFK immunoblot analysis with data shown as mean band densities normalized to total SFK (N=5/group). Quantitative real-time polymerase chain reaction (qRT-PCR) analysis for (**C**) *Cd36* (N=6-8/group); (**D**) *Il6* (N=5-7/group); (**E**) *Tnf* (N= 7/group); (**F**) *Ppargc1a* (N=6-7/group). For all qRT-PCR, GAPDH was used as a housekeeping gene. Values represent means ± SEM. *p<0.05 vs. *Apoe*^-/-^ chow; **p<0.01 vs. *Apoe*^-/-^ chow; ***p<0.001 vs. *Apoe*^-/-^ chow; ****p<0.0001 vs. *Apoe*^-/-^ chow; ^#^p<0.05 vs. *Apoe*^-/-^ WD; ^##^p<0.01 vs. *Apoe*^-/-^ WD; ^###^p<0.001 vs. *Apoe*^-/-^ WD; ^####^p<0.0001 vs. *Apoe*^-/-^ WD.

### Adipocyte-specific NaKtide expression improved glucose tolerance and systemic inflammation, and reduced expression of circulatory pro-inflammatory miRs in *Apoe*^-/-^ mice fed WD

Blood glucose levels in WD-fed *Apoe*^-/-^ mice were significantly elevated compared to chow-fed controls (Figure 5A), and Adipo-NaKtide treatment reduced these levels and improved glucose tolerance. Similarly, while WD increased plasma IL-6 and TNF-α levels to 297.46±10.20 pg/ml and 9.12±0.47 pg/ml, respectively, Adipo-NaKtide treatment lowered both to levels near the chow-fed controls (129.15±16.54 pg/ml for IL-6 and 7.36±0.44 pg/ml for TNF-α) (Figure 5B and 5C). Additionally, relative expression levels of circulating miR-27 b-3p and miR-34a-3p, known mediators of atherosclerosis development,^28,29^ were downregulated by Adipo-NaKtide treatment in WD diet-fed *Apoe*^-/-^ mice (Figure 5D and 5E).

**Figure 5.**
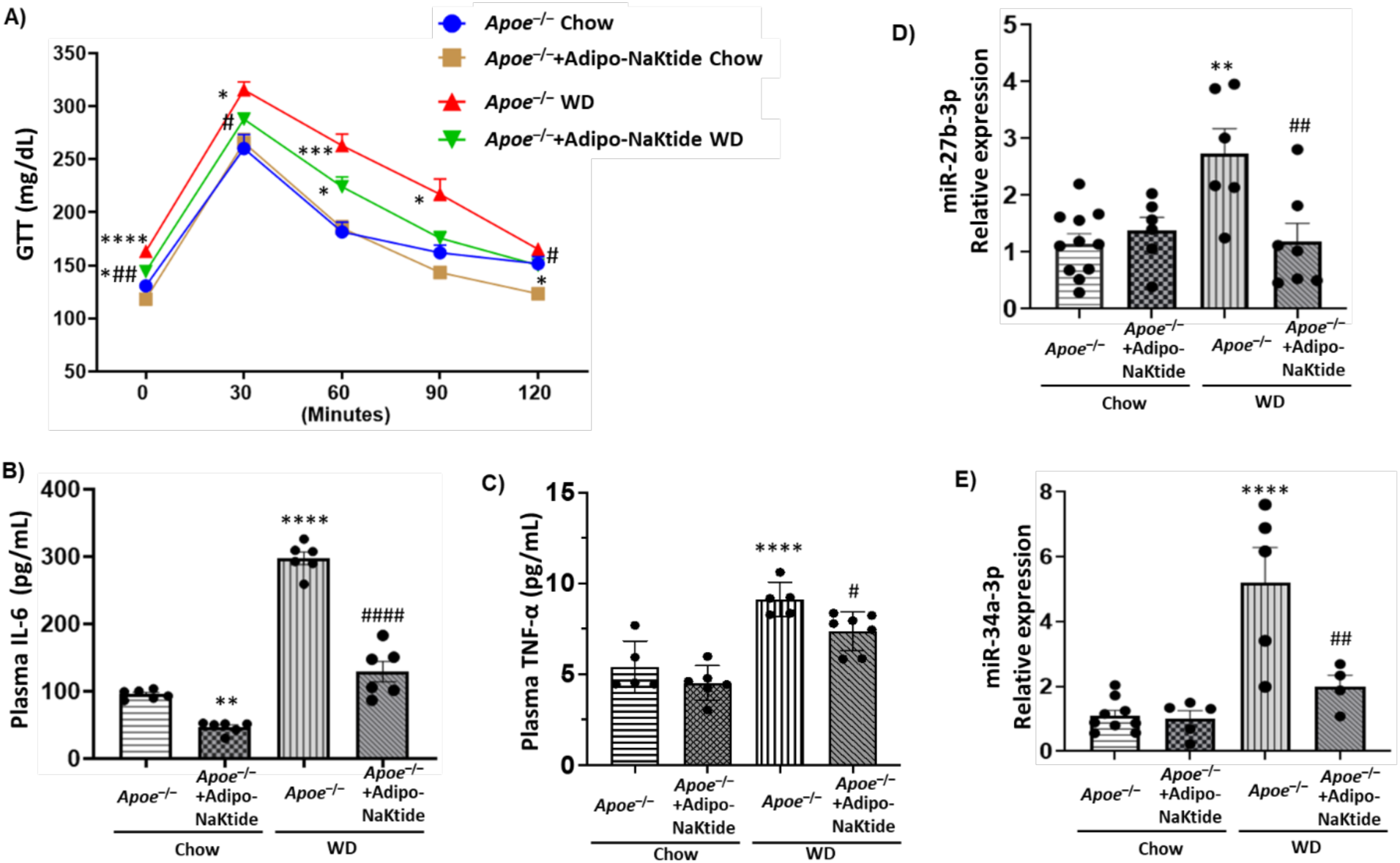
Adipocyte-specific expression of NaKtide improves metabolic profile, systemic inflammation, and circulating miRs in *Apoe*^-/-^ mice fed a WD. (**A**) Glucose tolerance test (N=9-10/group). Plasma concentration of (**B**) IL-6 (N=6/group); (**C**) TNF-α (N=5-7/group). Quantitative real-time polymerase chain reaction (qRT-PCR) analysis for (**D**) miR-27b-3p (N= 6-11/group); (**E**) miR-34a-3p (N= 4-9/group). For all qRT-PCR, U6 was used as an endogenous control. Values represent means ± SEM. *p<0.05 vs. *Apoe*^-/-^ chow; **p<0.01 vs. *Apoe*^-/-^ chow; ***p<0.001 vs. *Apoe*^-/-^ chow, ****p<0.0001 vs. *Apoe*^-/-^ chow; ^#^p<0.05 vs. *Apoe*^-/-^ WD; ^##^p<0.01 vs. *Apoe*^-/-^ WD; ^####^p<0.0001 vs. *Apoe*^-/-^ WD.

## DISCUSSION

Atherosclerosis is increasingly recognized as a disease influenced by systemic metabolic and inflammatory signals beyond the vasculature^30^. Adipose tissue has emerged as a key modulator of vascular health through its endocrine and paracrine functions^31^. While adipocyte dysfunction is known to exacerbate oxidative stress and inflammation^6,32^, the upstream signaling pathways mediating these effects remain incompletely understood. Here, we identify NKA signaling in adipocytes as a previously underappreciated contributor to atherogenesis.

Our study demonstrates that disruption of adipocyte NKA signaling via Adipo-NaKtide markedly attenuates atherosclerotic plaque development in WD-fed *Apoe*^-/-^ mice (Figure 1). These protective effects were accompanied by reduced macrophage accumulation in the aortic sinus (Figure 2), pointing to the importance of adipocyte-derived signals in shaping the vascular immune landscape. Notably, while the concept of adipose tissue influencing vascular inflammation is not new, our work provides the first *in vivo* evidence that NKA signaling within adipocytes is a key upstream regulator of these pro-atherogenic processes.

Mechanistically, we showed that Adipo-NaKtide suppressed adipocyte oxidative stress and SFK activation (Figure 4), a hallmark of NKA signal transduction^10^. This is consistent with our prior studies showing that NKA functions as a ROS amplifier via a feed-forward mechanism involving SFK activation and carbonylation of the NKA α1 subunit^33^. The suppression of pro-inflammatory cytokine expression and restoration of *Ppargc1a* levels (Figure 4) further suggest that adipocyte NKA signaling governs a broader metabolic-inflammatory program, likely through mitochondrial dysfunction and redox imbalance.

A novel aspect of our findings is the potential requirement of NKA signaling for CD36-mediated adipocyte activation. Our previous work established that CD36 and NKA form a signaling complex in adipocytes, and that oxLDL-induced CD36 signaling is blunted by pNaKtide^16^. Here, Adipo-NaKtide reduced *Cd36* transcript levels and abrogated SFK activation *in vivo* (Figure 4), supporting a model in which NKA is a key modulator or even a prerequisite for CD36-dependent pro-oxidant signaling in adipocytes. Given CD36’s established role in lipid uptake^34^, mitochondrial metabolism^20^, and immune signaling^35^, this NKA-CD36 axis may serve as a central regulatory node linking lipid handling to inflammatory output in adipose tissue.

Additionally, our data suggest that adipocyte NKA signaling influences systemic inflammation and glucose homeostasis (Figure 5). Adipo-NaKtide improved glucose tolerance and reduced circulating pro-inflammatory cytokines and miRs (miR-27b and miR-34a), both of which have been implicated in atherosclerosis^28,36^. These findings reinforce the concept of adipose tissue as an active participant in vascular disease and highlight NKA signaling as a modifiable upstream target.

It is intriguing that suppression of adipocyte NKA signaling also led to a reduction in α-SMA-positive cells in the aortic sinus, most likely smooth muscle cells (Figure 2). While the mechanisms remain unclear, this raises the possibility that adipocyte-derived factors regulated by NKA signaling can influence vascular smooth muscle phenotype and behavior in a non-cell-autonomous manner. Future studies are warranted to identify these factors and dissect their downstream effects.

We acknowledge several limitations. First, the identity of the specific adipocyte-derived mediators remains to be determined. Second, although our model achieves adipocyte-restricted NKA inhibition, the long-term effects and safety of targeting this pathway systemically remain to be established. Third, while our study provides strong evidence from a murine model, the relevance to human adipose tissue biology and atherosclerosis will require further validation.

In summary, we identify adipocyte NKA signaling as a novel driver of atherosclerosis, acting through oxidative stress amplification, modulation of inflammation, and potentially adipose-vascular communication (Figure 6). These findings establish a mechanistic framework for understanding how adipocytes contribute to vascular disease and open new avenues for exploring NKA signaling as a therapeutic target in metabolic and cardiovascular disorders.

**Figure 6.**
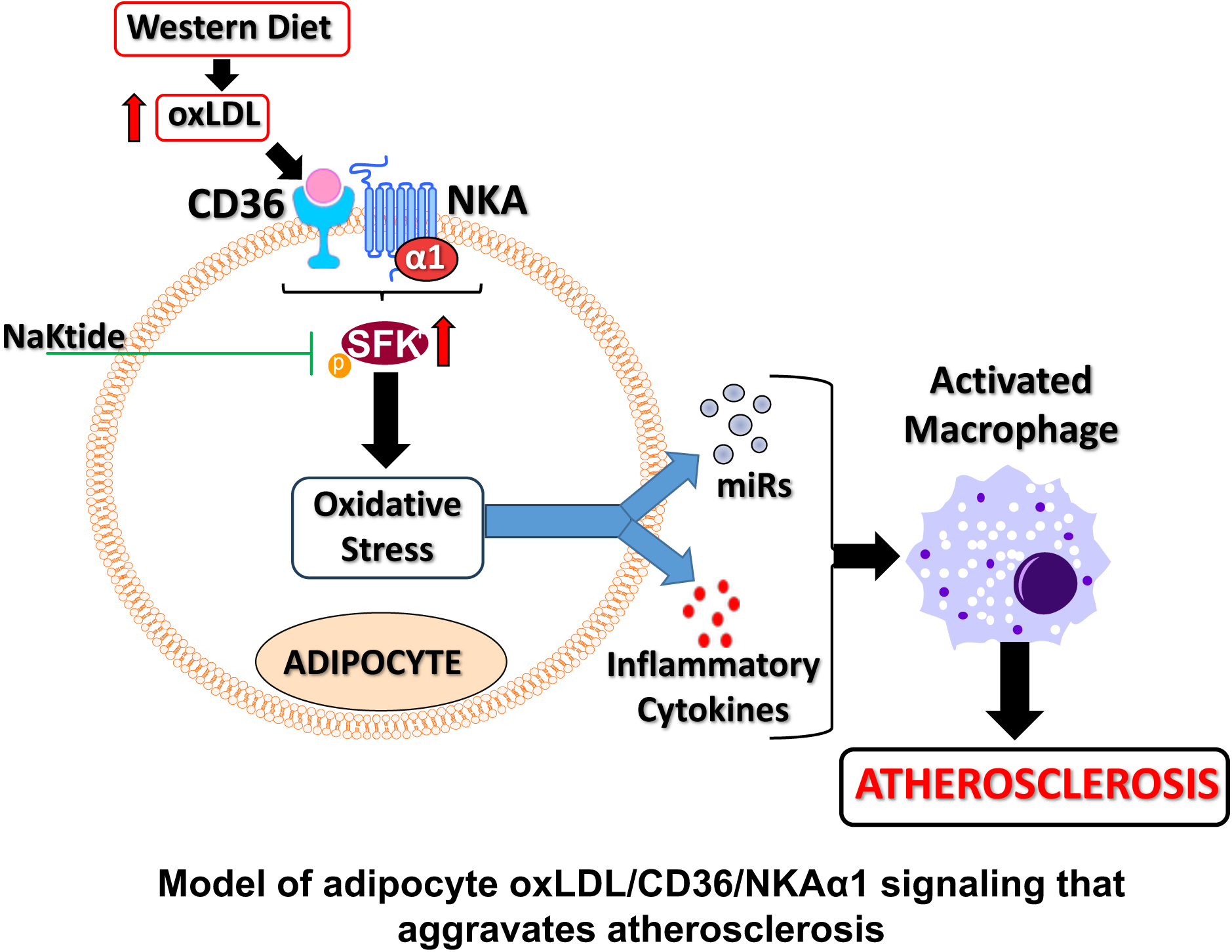
The schematic model of oxLDL/CD36/NKAα1 signaling that promotes crosstalk between adipocytes and macrophages aggravates atherosclerosis development.

## Acknowledgements

We thank Mr. Douglas Franklin and Ms. Vaya Chen for their excellent technical support. We also thank Versiti Blood Research Institute Shared Resources (RRID: SCR_025503) for their services, instrumentation, and specialist support.

## Supplemental Material

Supplemental Figure 1

## Authors Contributions

Bruno S. Goncalves and Yaxin Wang performed the experiments, wrote the manuscript, and analyzed data. Sneha S Pillai, Hari V. Lakhani, and Aslam Chaudhry performed the experiments and analyzed data. Roy L. Silverstein, Joseph I. Shapiro, Yiliang Chen and Komal Sodhi designed the experiments, interpreted the data, wrote and edited the manuscript. All authors contributed to the article and approved the submitted version.

## Data and materials availability

All data are available in the main text and supplemental materials.

## REFERENCES

1. Hulsmans M, Holvoet P. The vicious circle between oxidative stress and inflammation in atherosclerosis. J Cell Mol Med. 2010;14:70–78. doi: 10.1111/j.1582-4934.2009.00978.x

2. Kattoor AJ, Pothineni NVK, Palagiri D, Mehta JL. Oxidative Stress in Atherosclerosis. Curr Atheroscler Rep. 2017;19:42. doi: 10.1007/s11883-017-0678-6

3. Aboonabi A, Meyer RR, Singh I. The association between metabolic syndrome components and the development of atherosclerosis. J Hum Hypertens. 2019;33:844–855. doi: 10.1038/s41371-019-0273-0

4. Fantuzzi G, Mazzone T. Adipose tissue and atherosclerosis: exploring the connection. Arterioscler Thromb Vasc Biol. 2007;27:996–1003. doi: 10.1161/ATVBAHA.106.131755

5. van Dam AD, Boon MR, Berbee JFP, Rensen PCN, van Harmelen V. Targeting white, brown and perivascular adipose tissue in atherosclerosis development. Eur J Pharmacol. 2017;816:82–92. doi: 10.1016/j.ejphar.2017.03.051

6. Manna P, Jain SK. Obesity, Oxidative Stress, Adipose Tissue Dysfunction, and the Associated Health Risks: Causes and Therapeutic Strategies. Metab Syndr Relat Disord. 2015;13:423–444. doi: 10.1089/met.2015.0095

7. Furukawa S, Fujita T, Shimabukuro M, Iwaki M, Yamada Y, Nakajima Y, Nakayama O, Makishima M, Matsuda M, Shimomura I. Increased oxidative stress in obesity and its impact on metabolic syndrome. J Clin Invest. 2004;114:1752–1761. doi: 10.1172/JCI21625

8. Haas M, Askari A, Xie Z. Involvement of Src and epidermal growth factor receptor in the signal-transducing function of Na+/K+-ATPase. J Biol Chem. 2000;275:27832–27837. doi: 10.1074/jbc.M002951200

9. Haas M, Wang H, Tian J, Xie Z. Src-mediated inter-receptor cross-talk between the Na+/K+-ATPase and the epidermal growth factor receptor relays the signal from ouabain to mitogen-activated protein kinases. J Biol Chem. 2002;277:18694–18702. doi: 10.1074/jbc.M111357200

10. Xie Z, Askari A. Na(+)/K(+)-ATPase as a signal transducer. Eur J Biochem. 2002;269:2434–2439. doi: 10.1046/j.1432-1033.2002.02910.x

11. Tian J, Cai T, Yuan Z, Wang H, Liu L, Haas M, Maksimova E, Huang XY, Xie ZJ. Binding of Src to Na+/K+-ATPase forms a functional signaling complex. Mol Biol Cell. 2006;17:317–326. doi: 10.1091/mbc.e05-08-0735

12. Li Z, Cai T, Tian J, Xie JX, Zhao X, Liu L, Shapiro JI, Xie Z. NaKtide, a Na/K-ATPase-derived peptide Src inhibitor, antagonizes ouabain-activated signal transduction in cultured cells. J Biol Chem. 2009;284:21066–21076. doi: 10.1074/jbc.M109.013821

13. Sodhi K, Maxwell K, Yan Y, Liu J, Chaudhry MA, Xie Z, Shapiro JI. pNaKtide Inhibits Na/K-ATPase Signaling and Attenuates Obesity. J Clin Med Sci. 2023;7.

14. Liu J, Tian J, Chaudhry M, Maxwell K, Yan Y, Wang X, Shah PT, Khawaja AA, Martin R, Robinette TJ, et al. Attenuation of Na/K-ATPase Mediated Oxidant Amplification with pNaKtide Ameliorates Experimental Uremic Cardiomyopathy. Sci Rep. 2016;6:34592. doi: 10.1038/srep34592

15. Sodhi K, Srikanthan K, Goguet-Rubio P, Nichols A, Nawab A, Shah P, Chaudhry M, El-Hamdani M, Xie Z, Shapiro J. Inhibition of Na/K-ATPase signaling Attenuates Steatohepatitis and Atherosclerosis in Mice Fed a Western Diet. Cell Mol Biol (Noisy-le-grand). 2023;69:162–171. doi: 10.14715/cmb/2023.69.2.27

16. Pillai SS, Pereira DG, Zhang J, Huang W, Beg MA, Knaack DA, de Souza Goncalves B, Sahoo D, Silverstein RL, Shapiro JI, et al. Contribution of adipocyte Na/K-ATPase alpha1/CD36 signaling induced exosome secretion in response to oxidized LDL. Front Cardiovasc Med. 2023;10:1046495. doi: 10.3389/fcvm.2023.1046495

17. Silverstein RL, Febbraio M. CD36 and atherosclerosis. Curr Opin Lipidol. 2000;11:483–491. doi: 10.1097/00041433-200010000-00006

18. Kennedy DJ, Chen Y, Huang W, Viterna J, Liu J, Westfall K, Tian J, Bartlett DJ, Tang WH, Xie Z, et al. CD36 and Na/K-ATPase-alpha1 form a proinflammatory signaling loop in kidney. Hypertension. 2013;61:216–224. doi: 10.1161/HYPERTENSIONAHA.112.198770

19. Chen Y, Kennedy DJ, Ramakrishnan DP, Yang M, Huang W, Li Z, Xie Z, Chadwick AC, Sahoo D, Silverstein RL. Oxidized LDL-bound CD36 recruits an Na(+)/K(+)-ATPase-Lyn complex in macrophages that promotes atherosclerosis. Sci Signal. 2015;8:ra91. doi: 10.1126/scisignal.aaa9623

20. Chen Y, Yang M, Huang W, Chen W, Zhao Y, Schulte ML, Volberding P, Gerbec Z, Zimmermann MT, Zeighami A, et al. Mitochondrial Metabolic Reprogramming by CD36 Signaling Drives Macrophage Inflammatory Responses. Circ Res. 2019;125:1087–1102. doi: 10.1161/CIRCRESAHA.119.315833

21. Sodhi K, Wang X, Chaudhary MA, Lakhani HV, Zehra M, Nawab A, Cottrill CL, Bai F, Liu J, Sanabria JR, et al. Adipocyte Na, K-ATPase Signaling Attenuates Experimental Uremic Cardiomyopathy. Cell Mol Biol (Noisy-le-grand). 2023;69:197–206. doi: 10.14715/cmb/2023.69.5.31

22. Moore KJ, Tabas I. Macrophages in the pathogenesis of atherosclerosis. Cell. 2011;145:341–355. doi: 10.1016/j.cell.2011.04.005

23. Ross R, Glomset JA. Atherosclerosis and the arterial smooth muscle cell: Proliferation of smooth muscle is a key event in the genesis of the lesions of atherosclerosis. Science. 1973;180:1332–1339. doi: 10.1126/science.180.4093.1332

24. Aherrahrou R, Lue D, Perry RN, Aberra YT, Khan MD, Soh JY, Ord T, Singha P, Yang Q, Gilani H, et al. Genetic Regulation of SMC Gene Expression and Splicing Predict Causal CAD Genes. Circ Res. 2023;132:323–338. doi: 10.1161/CIRCRESAHA.122.321586

25. Cochain C, Vafadarnejad E, Arampatzi P, Pelisek J, Winkels H, Ley K, Wolf D, Saliba AE, Zernecke A. Single-Cell RNA-Seq Reveals the Transcriptional Landscape and Heterogeneity of Aortic Macrophages in Murine Atherosclerosis. Circ Res. 2018;122:1661–1674. doi: 10.1161/CIRCRESAHA.117.312509

26. Shirai T, Nazarewicz RR, Wallis BB, Yanes RE, Watanabe R, Hilhorst M, Tian L, Harrison DG, Giacomini JC, Assimes TL, et al. The glycolytic enzyme PKM2 bridges metabolic and inflammatory dysfunction in coronary artery disease. J Exp Med. 2016;213:337–354. doi: 10.1084/jem.20150900

27. Rius-Perez S, Torres-Cuevas I, Millan I, Ortega AL, Perez S. PGC-1alpha, Inflammation, and Oxidative Stress: An Integrative View in Metabolism. Oxid Med Cell Longev. 2020;2020:1452696. doi: 10.1155/2020/1452696

28. Tang Y, Yang LJ, Liu H, Song YJ, Yang QQ, Liu Y, Qian SW, Tang QQ. Exosomal miR-27b-3p secreted by visceral adipocytes contributes to endothelial inflammation and atherogenesis. Cell Rep. 2023;42:111948. doi: 10.1016/j.celrep.2022.111948

29. Pan Y, Hui X, Hoo RLC, Ye D, Chan CYC, Feng T, Wang Y, Lam KSL, Xu A. Adipocyte-secreted exosomal microRNA-34a inhibits M2 macrophage polarization to promote obesity-induced adipose inflammation. J Clin Invest. 2019;129:834–849. doi: 10.1172/JCI123069

30. Madaudo C, Coppola G, Parlati ALM, Corrado E. Discovering Inflammation in Atherosclerosis: Insights from Pathogenic Pathways to Clinical Practice. Int J Mol Sci. 2024;25. doi: 10.3390/ijms25116016

31. Mohamed-Ali V, Pinkney JH, Coppack SW. Adipose tissue as an endocrine and paracrine organ. Int J Obes Relat Metab Disord. 1998;22:1145–1158. doi: 10.1038/sj.ijo.0800770

32. Berg AH, Scherer PE. Adipose tissue, inflammation, and cardiovascular disease. Circ Res. 2005;96:939–949. doi: 10.1161/01.RES.0000163635.62927.34

33. Yan Y, Shapiro AP, Haller S, Katragadda V, Liu L, Tian J, Basrur V, Malhotra D, Xie ZJ, Abraham NG, et al. Involvement of reactive oxygen species in a feed-forward mechanism of Na/K-ATPase-mediated signaling transduction. J Biol Chem. 2013;288:34249–34258. doi: 10.1074/jbc.M113.461020

34. Hao JW, Wang J, Guo H, Zhao YY, Sun HH, Li YF, Lai XY, Zhao N, Wang X, Xie C, et al. CD36 facilitates fatty acid uptake by dynamic palmitoylation-regulated endocytosis. Nat Commun. 2020;11:4765. doi: 10.1038/s41467-020-18565-8

35. Kennedy DJ, Kuchibhotla S, Westfall KM, Silverstein RL, Morton RE, Febbraio M. A CD36-dependent pathway enhances macrophage and adipose tissue inflammation and impairs insulin signalling. Cardiovasc Res. 2011;89:604–613. doi: 10.1093/cvr/cvq360

36. Su G, Sun G, Liu H, Shu L, Liang Z. Downregulation of miR-34a promotes endothelial cell growth and suppresses apoptosis in atherosclerosis by regulating Bcl-2. Heart Vessels. 2018;33:1185–1194. doi: 10.1007/s00380-018-1169-6

